# Discovery of another mechanism for the inhibition of particulate guanylyl cyclases by the natriuretic peptide clearance receptor

**DOI:** 10.1101/2022.09.06.506848

**Authors:** Dianxin Liu, Ryan P. Ceddia, Wei Zhang, Fubiao Shi, Huafeng Fang, Sheila Collins

## Abstract

The cardiac natriuretic peptides (NPs) control pivotal physiological actions such as fluid and electrolyte balance, cardiovascular homeostasis, and adipose tissue metabolism by activating their receptor enzymes (NPRA and NPRB). These receptors are homodimers that generate intracellular cyclic guanosine monophosphate (cGMP). The NP receptor NPRC, nicknamed the clearance receptor, lacks a guanylyl cyclase domain; instead, it can bind the NPs to internalize and degrade them. The conventional paradigm is that by competing for and internalizing NPs, NPRC blunts the ability of NPs to signal through NPRA and NPRB. Here we show another previously unknown mechanism by which NPRC can interfere with the cGMP signaling function of the NP receptors. By forming a heterodimer with monomeric NPRA or NPRB, NPRC can prevent the formation of a functional guanylyl cyclase domain and thereby suppress cGMP production in a cell-autonomous manner.

**Significance Statement:** Natriuretic peptides (NP) are hormones that are established regulators of vascular and cardiac function, in part through their regulation of fluid and electrolyte balance. NPs signal through particulate guanylyl cyclases (NPRA and NPRB), which are homodimeric membrane-bound receptor enzymes that generate cGMP upon NP binding. Additionally, a ‘silent’ NP receptor (NPRC) lacks the guanylyl cyclase domain and is a negative regulator of NP signaling. It has been demonstrated that NPRC undergoes internalization and recycling and thus removes NPs, thereby blunting activation of the guanylyl cyclase-containing receptors. Here we show an additional mechanism by which NPRC inhibits NP signaling. Our results show that NPRC can directly interact with NPRA and NPRB, forming non-functional receptor heterodimers with NPRA and NPRB, thereby abrogating NP-evoked cGMP production. This finding establishes another novel mechanistic role for NPRC.

## Introduction

Following the unexpected finding that atrial cells contained ‘granules’ reminiscent of secretory tissues (1), in 1981 de Bold and colleagues (2) made the observation that when extracts from atria were injected into rats it caused a major increase in natriuresis and diuresis and a significant drop in blood pressure. These natriuretic factors were then shown to be peptides (3-8). We now know these factors as atrial natriuretic peptide (ANP) and B-type natriuretic peptide (BNP). These discoveries unexpectedly placed the heart in the category of an endocrine organ. It was subsequently determined that the action of these natriuretic factors was mediated by a receptor, natriuretic peptide receptor-A (NPRA, also referred to as GC-A), in the kidney to cause natriuresis and diuresis. NPRA is an integral membrane protein that in dimeric form contains guanylyl cyclase activity (9) and the second messenger cyclic GMP (cGMP) mediates NP action. Another related peptide, CNP, was later discovered that has its own guanylyl cyclase-containing receptor NPRB (or GC-B) (10). A third NP receptor, NPRC, was discovered (11) and referred to as the ‘silent’ receptor (12), as it lacks guanylyl cyclase activity, has a short cytoplasmic domain, and removes NPs from the circulation by endocytosis. This was discovered when administration of variants of ANP_1-28_, including ANP_4-23_ and a modified ring-deleted form that is selective for binding NPRC (12, 13), were found to increase endogenous levels of ANP and BNP in circulation (12-15), suggesting it played some role in antagonizing the actions of the GC-containing receptors. NPRC-mediated endocytosis of ANP was observed in several subsequent studies (16-19), further cementing NPRC’s nickname as the ‘NP clearance receptor’. As a result, NPRC can be considered to function as an inhibitor of NPRA/NPRB-cGMP signaling by competing for the peptide ligands. For homodimeric NPRA and NPRB, whether they internalize and recycle is unclear, as studies using different models have led to inconsistent results (17, 20-26).

Since the initial discovery of the cardiovascular effects of NPs, they have been found to exert a broader range of physiological functions beyond natriuresis and vasodilation, ranging from protection from cardiac fibrosis to stimulating adipose, bone, and skeletal muscle metabolism (27, 28). Clinically, the level of NPRC in tissues has been correlated with certain disease states. For example, in large population studies, subjects with obesity and hypertension have been shown to have lower circulating ANP and BNP levels (29-35), coupled with a dramatic increase in the level of NPRC in their adipose tissue (36-40). This has led to the concept of a ‘natriuretic handicap’, and it has been proposed that possibly the increased adipose NPRC levels may serve as a ‘sink’ to remove NPs from the circulation and thus contribute to disease pathology in terms of blunted adipose tissue lipolysis as well as increased blood pressure and risk of heart failure (30). This increase in NPRC levels with obesity has also been observed in preclinical models, including rats and mice (41-43). Conversely, in both rodents and humans, dieting/fasting and cold exposure raises the NPRA/NPRC ratio in adipose tissue, largely by decreasing the levels of NPRC, resulting in improved NP signaling (44-46). In support of this we recently reported that adipose tissue-specific disruption of the *Nprc* gene in mice resulted in improvements in several aspects of metabolic disease in an obesity paradigm (43). As we and others showed in earlier work, raising the level of NPRC shifts the dose-response curve for cGMP production through NPRA to the right in the same manner as a receptor antagonist (43, 47). Thus, the biological significance of the relative ratio of NPRA to NPRC *in vivo* raises questions such as how NPRC expression is regulated at the transcriptional level. Recently we and others have begun to address this issue (48-51). In this study, we make the observation that elevation of NPRC in cells appears to be correlated with lower NPRA protein levels. In addition to its clearance function, in the present study we investigated whether NPRC might also form heterodimers with NPRA or NPRB, thus preventing cGMP production. In this report we present evidence for the existence of such heterodimers using several independent approaches.

## Results

As is known from earlier studies (41), the NP receptors are expressed in a wide variety of tissues (**Figure 1A**). Numerous reports have shown that NPRC opposes NP-evoked signaling (27, 43, 52, 53) with the current understanding being that NPRC does not affect NPRA/B *per se*, but rather binds and removes the ligands. NPRC is well known to be endocytosed in a clathrin-mediated manner (16-19), and it is thought that this endocytosis and recycling reduces the local concentration of NPs. Interestingly, when NPRC levels are increased in adipose tissue through the well-known mechanism of high fat diet (HFD) induced obesity (37, 42, 51, 54, 55), we observe a modest decrease in the amount of NPRA on the plasma membrane in wild-type animals but not in those with a deletion of NPRC (**Figure 1B**). This did not appear to be due to NPRA protein moving to the cytosolic fraction as cytosolic NPRA remained unchanged by HFD feeding (**Figure 1C**). When we transfected the mouse brown adipocyte cell line, HIB-1B, with a FLAG-NPRA containing plasmid, we observed an interesting effect of increased NPRA protein upon proteosome inhibition (**Figure 1D**). The Flag-NPRA was moderately expressed in HIB-1B adipocytes and was further reduced by overexpression of exogeneous NPRC. This led us to speculate that NPRC was somehow suppressing the levels of Flag-NPRA protein. Treating the cells with the proteosome inhibitor MG-132 greatly increased Flag-NPRA protein levels in cells without exogeneous NPRC, and in cells with exogeneous NPRC, MG-132 partially restored Flag-NPRA levels. This observation may be somewhat cell type specific as other work in this manuscript shows that this plasmid construct and other NPRA expressing plasmids work well in HEK-293 cell lines that also express a moderate amount of NPRC. Overall, **Figure 1** shows an interesting correlation between high NPRC expression and low levels of NPRA on the plasma membrane when expressed in the same cell. From these findings in **Figure 1**, we cannot imply any causal link between NPRA and NPRC protein levels, but we thought that this subject deserved further investigation, so we conducted a series of studies to determine how NPRC could affect NPRA in a cell-autonomous manner.

**Figure 1.**
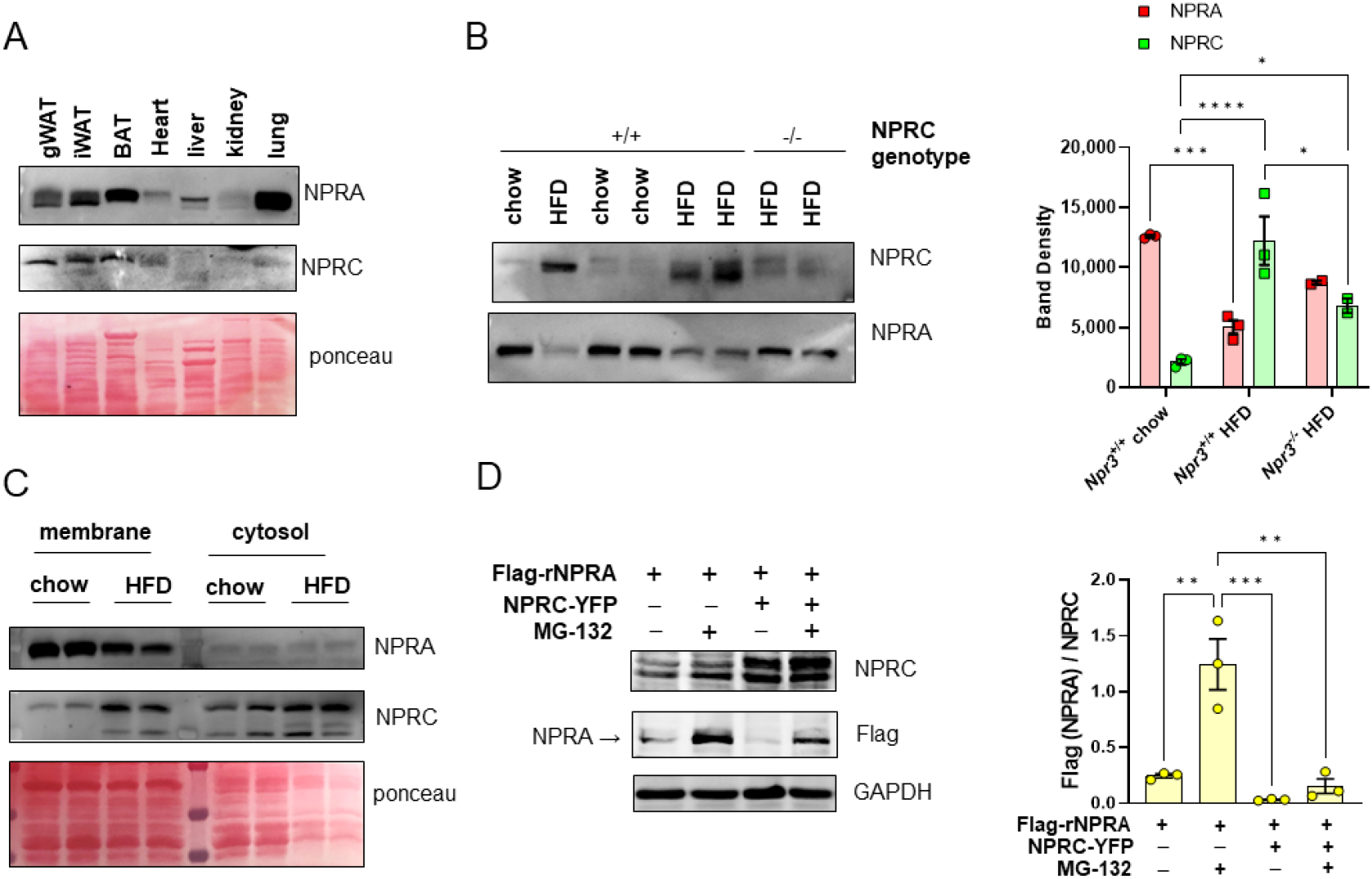
Comparison of NPRA and NPRC protein expression. (**A**) Protein expression of NPRA and NPRC were visualized by Western Blotting in membrane enriched fractions prepared from a variety of mouse tissues: gonadal white adipose tissue (gWAT), inguinal white adipose tissue (iWAT), brown adipose tissue (BAT), heart, liver, kidney, and lung. (**B**) Brown Adipose Tissue (BAT) membrane enriched fractions from wild type and NPRC^-/-^ mice fed either standard chow or high fat diet (HFD) were used to visualize NPRA and NPRC protein levels. Šídák’s multiple comparisons test for two-way ANOVA indicated on graph. (**C**) BAT protein from chow and HFD fed mice was separated into membrane and cytosolic enriched fractions. NPRA and NPRC were assessed by Western Blotting. (**D**) HIB-1B cells were transfected with the indicated plasmids and treated with adipocyte differentiation cocktail for 48-hours. Cells were then treated with the 2.5 µM MG-132 for 16 hours. Flag-NPRA and NPRC were assessed from whole cell lysates by Western Blotting. Šídák’s multiple comparisons test for one-way ANOVA indicated on graph.

Our first approach was to utilize co-immunoprecipitation to determine if NPRA/B could associate with NPRC. **Figure 2A** shows that following transfection of Flag-tagged NPRA or NPRB into HEK-293 cells, immunoprecipitation of the cell lysate with anti-Flag antibody retrieves the endogenously expressed NPRC protein while vascular endothelial growth factor receptor-1 (VEGFR1), an unrelated single trans-membrane receptor that also dimerizes, was unaffected. We attempted to use GC-C as a negative control, as it is also a particulate guanylyl cyclase that binds guanylin and uroguanylin instead of NPs; however, as can be seen in **Figure 2B**, GC-C also co-immunoprecipitated with NPRC, suggesting that NPRC may also interact with the particulate guanylyl cyclases that do not bind NPs. **Figure 2C** provide quantifications from independent experiments. **Figure 2D** shows that this heterodimer interaction is independent of ligand, whether using ANP_1-28_ or the NPRC-selective ANP_4-23_. In order to understand which cellular domain is responsible for this protein-protein interaction, a truncated form of NPRA was generated that retains the transmembrane domain but lacks all except 37 amino acids of the intracellular domain; thus making its intracellular domain the same length as NPRC (**Fig. S1A**). As shown in **Figure 2E**, similar to the full length NPRA, the truncated form could also immunoprecipitate the endogenous NPRC. Since the majority of the kinase homology domain and the entire hinge region and guanylyl cyclase catalytic domain were removed, it can be concluded that these are not necessary for heterodimer formation. In support of this a crystal structure of the dimeric extracellular hormone binding domain of NPRA (56) suggests that this region is sufficient for dimer formation, and Chinkers and Wilson previously showed that NPRA constructs containing only the extracellular domain, transmembrane and 5 amino acids of the intracellular domain were sufficient to retain homodimerization (57).

**Figure 2.**
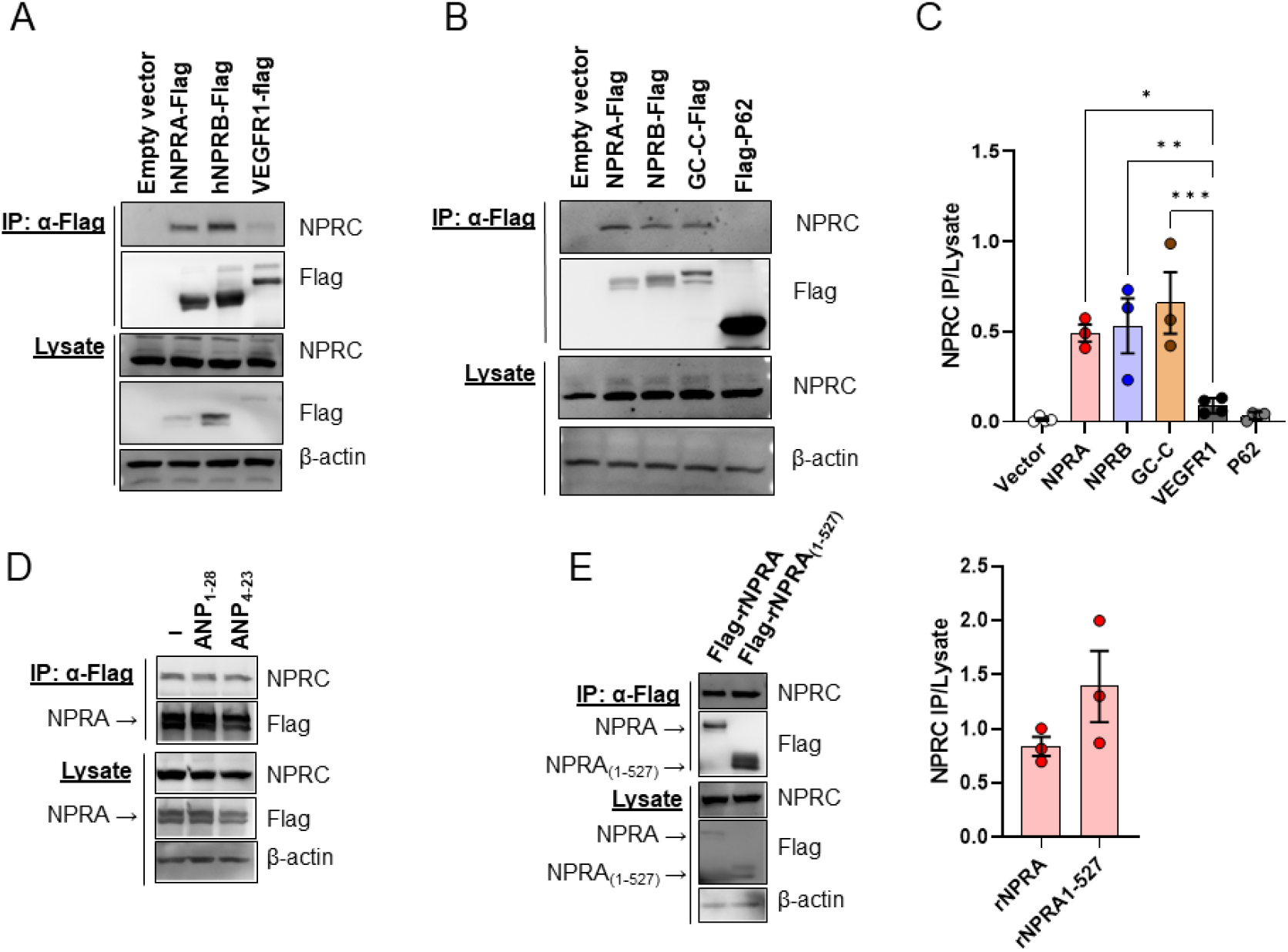
NPRC immunoprecipitates with NPRA and NPRB. HEK 293FT cells were transfected with plasmids expressing Flag-tagged natriuretic peptide receptors. Twenty-four hours after transfection, cell lysates were immunoprecipitated with anti-Flag Affinity agarose. (**A.**) Endogenous NPRC coimmunoprecipitated with hNPRA-Flag, hNPRB-Flag, or VEGFR1-Flag. NPRC band intensity to input of NPRC was calculated through Image J software was presented in the right panel. Šídák’s multiple comparisons test for one-way ANOVA comparing all samples to the negative control VEGFR1 for Western Blots in **A** and **B** are indicated on graph. (**B.**) Endogeneous NPRC coimmunoprecipitated with GC-C-Flag. (**C.**) hNPRA-Flag expressing HEK 293FT cells were treated with ANP (ANP_1-28_, 200 nM) or a truncated form of ANP that preferentially binds NPRC (ANP_4-23_, 200 nM) for 30 minutes. Neither ANP_1-28_ or ANP_4-23_ affected the ability of endogenous NPRC to coimmunoprecipitate with hNPRA-Flag. (**D.**) Endogenous NPRC coimmunoprecipitated with Flag-rNPRA and a truncated version of Flag-rNPRA lacking the c-terminal amino acids 528-1057 (Flag-rNPRA_(1-527)_).

Our second approach was to use a complementation reporter assay, specifically the NanoLuc® Binary Technology (NanoBiT) (58), to study protein-protein interactions between NPRA/B and NPRC. This technique allows for the observation of heterodimers in living cells and avoids relying upon antibodies for their detection. We made NPRA, NPRB, and NPRC constructs that contain a ‘two-hybrid’ type luciferase reporter at their C-termini (see **Fig. S1B**). In this assay, the luciferase signal is only produced when proteins containing the complementary halves of the reporter subunits (Large BiT, *lg*, and Small BiT, *sm*) form a functional luciferase enzyme. For this assay HAP-1 cells were chosen due to low endogenous expression level of all three natriuretic peptide receptors (**Fig. 3**). **Figure 4A** shows that when NPRA and NPRC are tagged with *lg* and *sm* complementary halves of the reporter subunits, significant luminescence is observed. The relative expression of these proteins is shown in the Western Blots in **Figure 4B**. While the strength of the luciferase signals varied across constructs, this is likely due to the relative levels of expression of the proteins and the size of the intracellular domains that could increase the distance between the two parts of the luciferase enzyme. This is especially true for the NPRC-*lg*/NPRA-*sm* in **Figure 3A**, which was expected due to the presence in full-length NPRA of the intracellular kinase homology domain, hinge region, and guanylyl cyclase catalytic domain, for which there is a difference of 528 intracellular amino acids in length between them. Because results from others (56, 57) and our data here (**Fig. 2E**) demonstrated that the intracellular guanylyl cyclase domain is not required for dimer formation, in subsequent experiments of this type we used the luciferase reporters for NPRA and NPRB that lack all but 37 amino acids of the intracellular domain termed: NPRA_1-532_ and NPRB_1-515_. As shown in **Fig. 4A** and **4C**, NPRC formed heterodimers with both NPRA_1-532_ and NPRB_1-515_ producing a strong luciferase response. To determine if expressing NPRC is capable of disrupting NPRA or NPRB homodimers in this assay, cells expressing NPRA_1-532_ or NPRB_1-515_ were co-transfected with Flag-tagged-NPRC. As shown in **Figure 4E, F** the presence of NPRC significantly reduced the luciferase signal, indicating the blunted ability of NPRA and NPRB to form homodimers by forming heterodimers instead.

**Figure 3.**
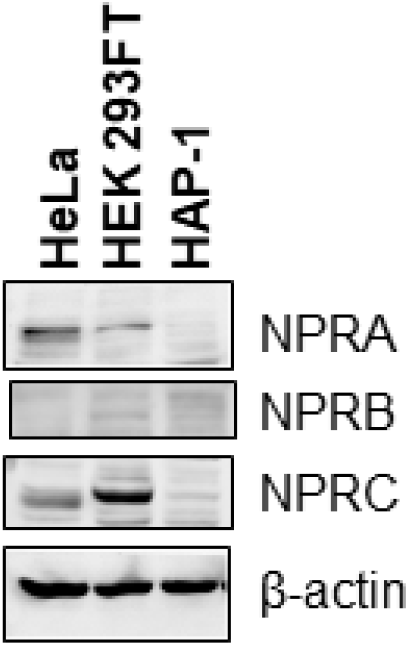
Endogenous natriuretic peptide receptor expression in cell models used. Expression of NPRA, NPRB, and NPRC were detected by Western Blot in HeLa, HEK293FT, and HAP-1 cells. β-actin was used as the loading control.

**Figure 4.**
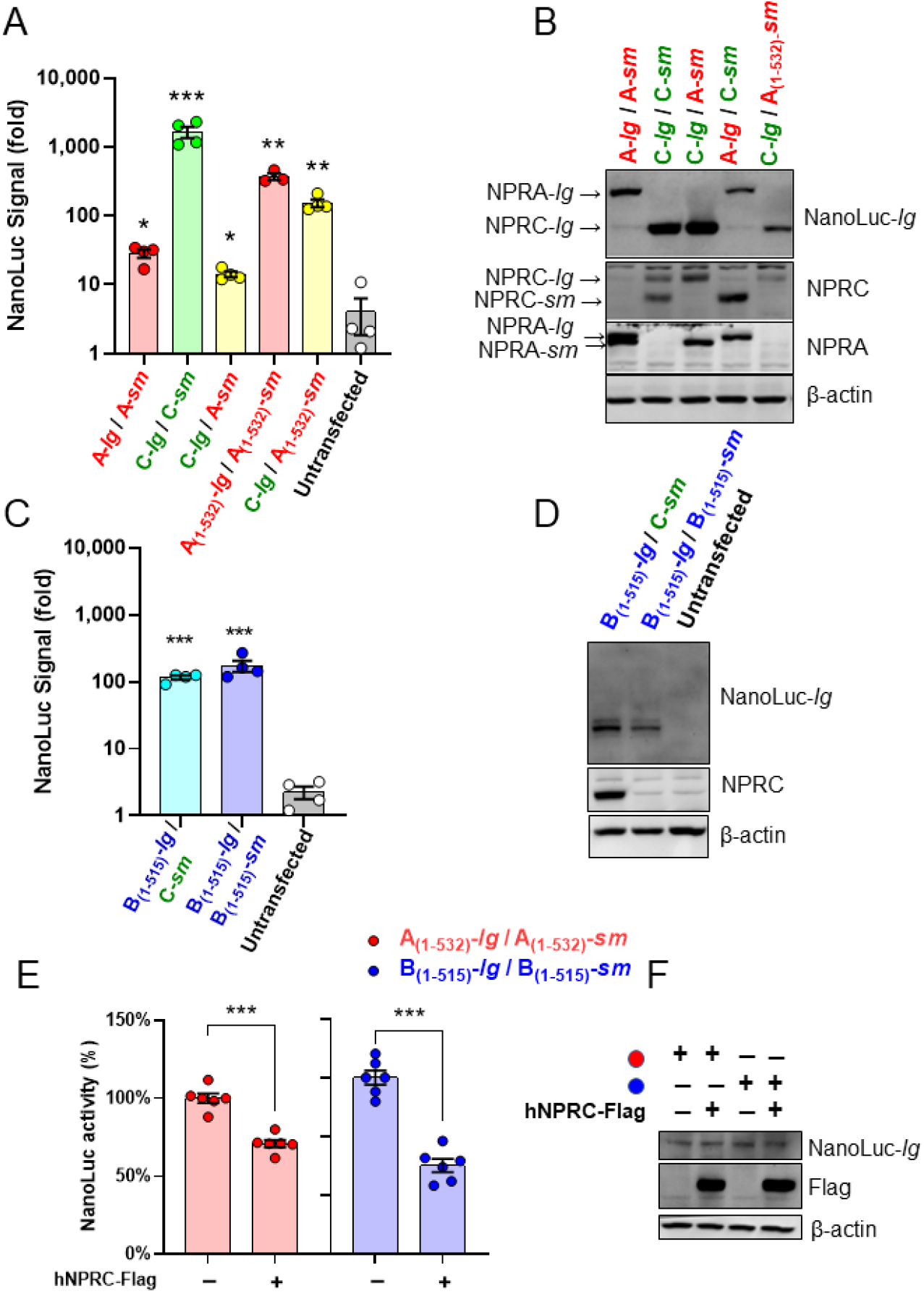
NPRC can form a heterodimer with either NPRA or NPRB and interferes with NPRA and NPRB homodimer formation. HAP-1 cells were transfected with natriuretic peptide receptors tagged with opposite halves of the NanoBiT complementation reporter. (**A**) A strong luciferase signal was observed for NPRA and NPRC homodimer as well as NPRA:NPRC heterodimer formation. A truncated version of NPRA containing only 37 intracellular amino acids (NPRA_(1-532)_) also formed homodimers and heterodimers with NPRC. (**B**) Transfected proteins were visualized by Western Blot. (**C**) The truncated version of NPRB containing only 37 intracellular amino acids (NPRA_(1-515)_) formed homodimers and heterodimers with NPRC. Full length NPRB exhibited poor expression, consequently the luciferase signal was not significantly greater than background (data not shown). (**D**) Western Blot was used to visualize expression of the transfected proteins. (**E**) HAP-1 cells were transfected with complementary halves of truncated NPRA or NPRB plus either unlabeled NPRC or empty vector. Both NPRA and NPRB expressing cells demonstrated reduced luciferase activity when expressing NPRC. (**F**) The transfected proteins were visualized by Western Blot. For all experiments, each point represents the average of 3 replicates from independent experiments. Paired t-tests were performed on the log transformed data comparing each treatment group to the untransfected control. * P<0.05, ** P<0.01, *** p<0.001.

Our third approach was to use a proximity ligation assay, specifically the DuoLink® proximity ligation assay (59), to study endogenous protein-protein interactions between NPRA and NPRC. We used HeLa and A-10 cells as models to test the NPRA and NPRC interaction because these cells endogenously express relatively high levels of both receptors (60). DuoLink® produces discreet fluorescent puncta when the antibodies for NPRA and NPRC are within 40 nm of each other. **Figure 5** shows that for both cell types, fluorescent puncta are detected only when antibodies for both receptors are present, again indicating that heterodimers of NPRA and NPRC can occur. Importantly, these results indicate that the endogenous receptors can form heterodimers under normal conditions.

**Figure 5.**
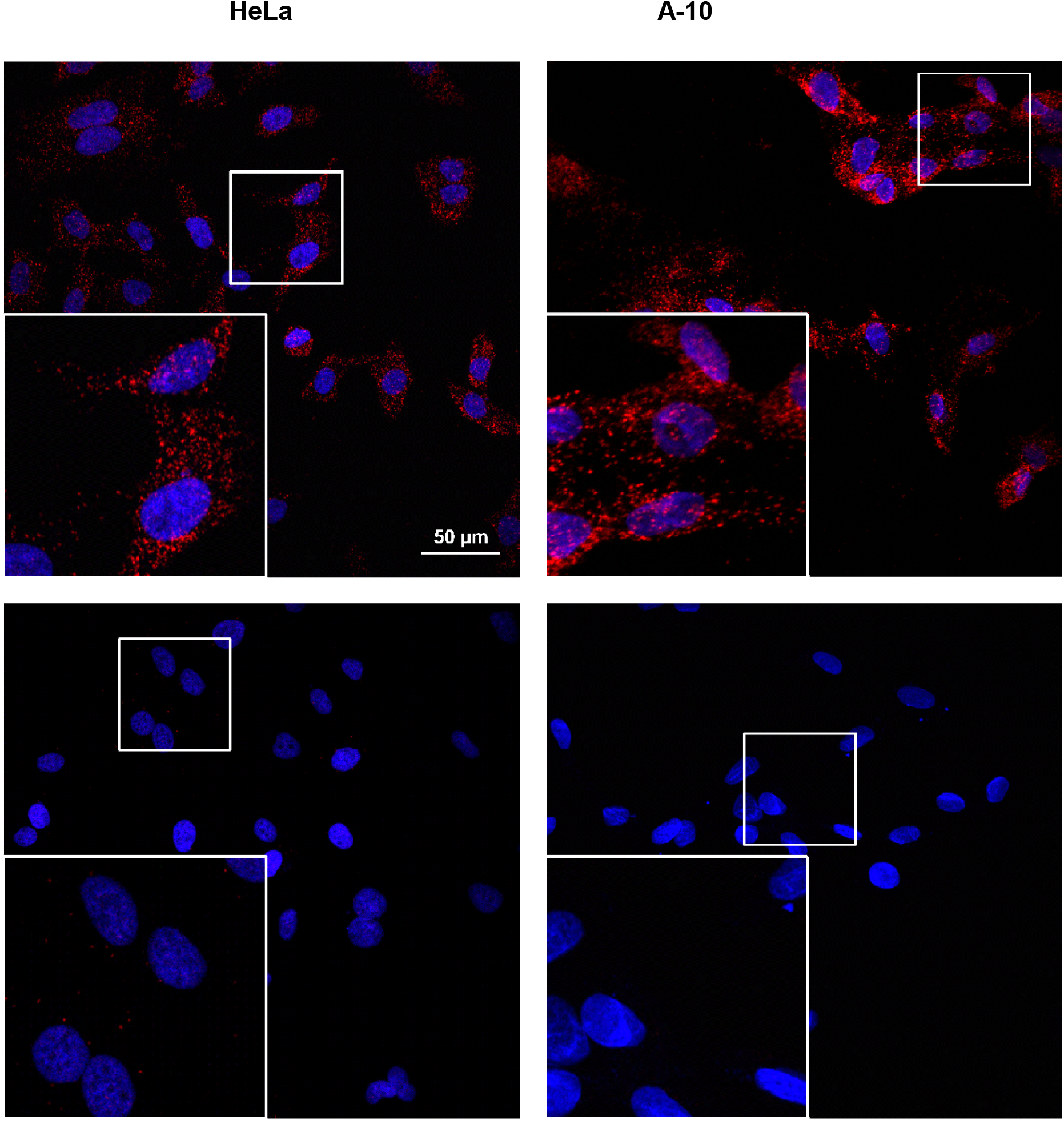
Endogenous NPRA:NPRC heterodimer formation in HeLa and A-10 cells. HeLa cells (left panels) and A-10 cells (right panels) were used in the Duolink Proximity Ligation Assay as described in the Methods with either NPRA and NPRC antibodies together (top) or NPRC antibody alone (bottom). Blue is DAPI stained nuclei and red is Duolink protein-protein interaction. Images are representative of at least three independent experiments.

These three approaches all indicate that heterodimers with NPRC can occur. We predict that the function of such heterodimers is to prevent cGMP generation because the heterodimer only contains one-half of the guanylyl cyclase enzyme, which is as an obligate homodimer (57, 61, 62). In order to examine the effects of NPRC on NPRA’s enzymatic activity, HEK-293 cells were transfected to express either NPRA alone or NPRA and NPRC together, followed by preparation of plasma membranes that were used in ANP-evoked cGMP assays. In this approach, unlike using intact cells as in our previous work (43), NPRC internalization to remove ANP cannot occur in a plasma membrane preparation. This greatly reduces, but does not eliminate, competition between NPRA and NPRC homodimers for binding ANP, but it does eliminate the internalization and recycling mechanism. As shown in **Figure 6** and **Fig. S2**, when NPRA and NPRC were expressed together in the same cell, the cGMP generated over time was significantly reduced, compared to plasma membranes from cells expressing NPRA alone. The Western Blots do not suggest differences in the amount of NPRA protein in these membranes leading us to conclude that the presence of NPRC itself reduces NPRA’s enzymatic activity presumably by the formation of catalytically inactive heterodimers. Therefore, if the effect of NPRC on NPRA function is solely due to ANP endocytosis and clearance, we would expect that there should be little to no difference in the rate of cGMP production in these plasma membrane preparations. Instead, this study supports the preceding data in that the formation of heterodimers with NPRC is responsible for a general decrease in NPRA activity.

**Figure 6.**
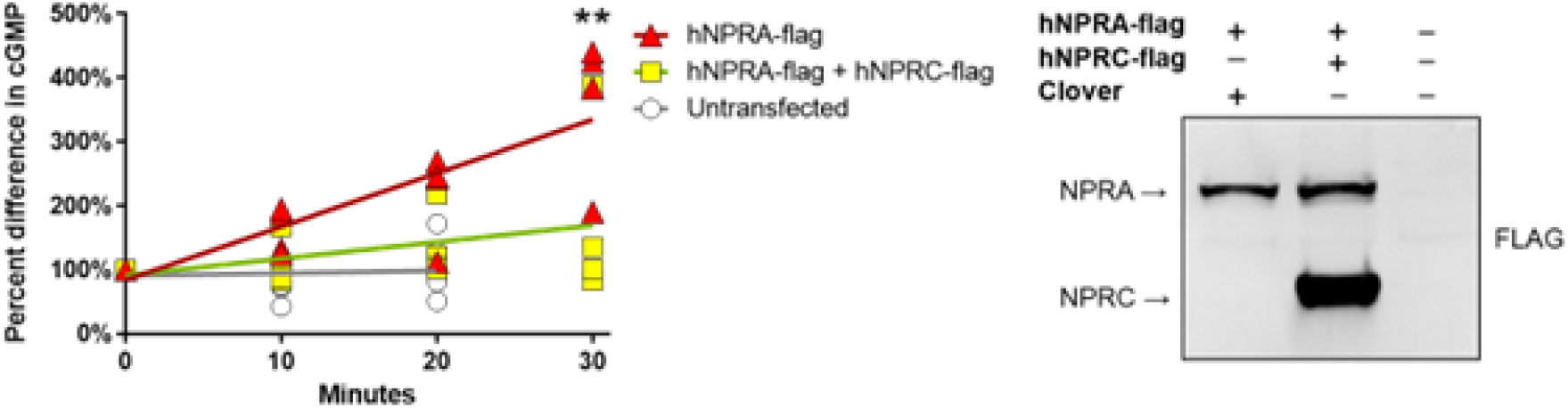
Co-expression of NPRC reduces ANP-evoked cGMP production in plasma membranes. Plasma membranes were made from HEK-293T cells transfected with either NPRA-flag plus pcDNA3-Clover control or NPRA-flag plus NPRC-flag. Plasma membranes were made without or with (**Fig S3**) phosphatase inhibitors. Co-expression of hNPRC along with hNPRA reduced the cGMP generated in the presence of 100 nM ANP (P=0.0163, test for difference in slopes between NPRA and NPRA+NPRC; Šídák’s multiple comparisons test comparing NPRA vs. NPRA+NPRC are indicated on the graphs). Untransfected cells did not appear to generate cGMP. Western Blot of membranes used in the ANP-stimulated cGMP assay. Co-expression of hNPRC along with hNPRA reduced the cGMP generated in the presence of 100 nM ANP (P=0.0203, test for difference in slopes between NPRA and NPRA+NPRC; Šídák’s multiple comparisons test comparing NPRA vs. NPRA+NPRC are indicated on the graphs). Cells transfected with pcDNA3-Clover did not appear to generate cGMP. Western Blot of membranes used in the ANP-stimulated cGMP assay.

## Discussion

In these studies, we identify a previously unknown mechanism by which NPRC can inhibit NP signaling. This discovery of the formation of non-functional heterodimers between NPRC and NPRA or NPRB does not challenge the veracity of the extensive prior work showing that NPRC can abrogate NP-evoked signaling by clearing the ligand, but rather demonstrates an additional mechanism for attenuating NP’s actions. This finding has profound implications for how we interpret NPRC’s opposition to NP signaling, as these studies reveal a novel cell autonomous mechanism for NPRC’s actions, which likely acts alongside the classical non-cell-autonomous endocytic and recycling mechanism. Because both the “clearance” and “heterodimerization” mechanisms are functional inhibitors of NP-evoked cGMP production and downstream signaling events, it is difficult to parse out the relative contribution of these mechanisms *in vivo* as tissues often express some mixture of NP receptors. What becomes more important to discern is which cell types within a tissue these receptors are expressed, since the heterodimer mechanism depends on the receptors being expressed within the same cell.

We have observed NPRC heterodimer formation with both transfected and endogenously expressed receptors and, in addition to techniques relying on protein-specific antibodies, these heterodimers have been detected using a complementation reporter assay that is not influenced by antibody fidelity. A limitation of this current work is that we have only been able to conclusively demonstrate the heterodimer formation in cell models. Nevertheless, understanding this heterodimerization process is important for interpreting studies utilizing co-expression or cell lines with high NPRA/B and NPRC levels. Currently, observing heterodimer formation in native tissue poses additional challenges due to the necessity for NPRA/B and NPRC to both be expressed in the same cell, physically near each other in the plasma membranes, for enough time to be captured, and in sufficient quantities to be detected. As many tissues do not endogenously express high levels of both receptors, the opportunities to observe heterodimers *in vivo* are limited though they may be occurring below detectable levels. This lack of high endogenous NPRA/B and NPRC co-expression may be associated with the controversy as to whether homodimeric NPRA can undergo clathrin-mediated internalization. While unproven, we speculate that NPRA may need to form a heterodimer with NPRC in order to be internalized. This is consistent with earlier observations that NPRA is most likely to internalize in cells that express a substantial fraction of NPRC (17). Prior to our discovery of NPRC-containing heterodimers, these observations appeared to be unconnected as NPRA and NPRC were thought to not interact directly. Our use of the proteosome inhibitor further suggests that it may be difficult to observe internalized NPRA if it is being rapidly degraded while NPRC is being recycled back to the plasma membrane. If future studies validate this hypothetical model, it will support our observed correlation between high NPRC and low NPRA protein expression.

All together our findings demonstrate that NPRC can directly interact with both NPRA and NPRB and competes with NPRA and NPRB for homodimer formation, with the net effect to reduce NP receptor evoked cGMP production. Our data indicate that in addition to its role as a clearance receptor, a previously unrecognized function of NPRC is to act as a dominant-negative NP receptor by forming catalytically inactive heterodimers with NPRA and NPRB. The consequence of shifting the proportions of NPRA (and NPRB) away from the homodimeric state and towards the heterodimeric state with NPRC is a reduced capacity to generate cGMP. We suggest that in addition to the non-cell autonomous mechanism of NPRC to dampen NP-evoked cGMP production by ligand clearance, this heterodimer formation may be a previously unidentified cell autonomous mechanism by which NPRC inhibits NP-cGMP signaling.

## Materials and Methods

### Plasmids

A summary of the plasmids used in this study can be found in **Supplemental Table 1**. NanoLuc plasmids were made by cloning full-length and truncated human *NPR1*, *NPR2*, and *NPR3* genes from the FLAG-tagged plasmids purchased from GenScript into the appropriate vectors from the NanoBiT® PPI MCS Starter System (Promega, #N2014). We also generated a rat NPRA plasmid lacking most the intracellular domain (Flag-rNPRA_(1-527)_), which was performed by site-directed mutagenesis to introduce an early stop codon and all plasmid constructs were sequenced and confirmed by Genewiz (South Plainfield, NJ). Primers used for cloning and sequencing can be found in **Supplemental Table 2**.

### Reagents and Antibodies

The following reagents were purchased from Sigma: 6-aminohexanoic acid (#A2504), bovine serum albumin (BSA) (#A7906), creatine phosphokinase (#C3755), digitonin (#D141), imidazole (#I202), guanosine 5′-triphosphate sodium salt (GTP) (#G8877), hydrogen peroxide solution (#216763) phenylmethanesulfonyl fluoride (PMSF) (#P7626), phosphocreatine disodium salt (#P7936). MG-132 was from Tocris (#1748). ANP_1-28_ was from AnaSpec (#AS-20648). ANP_4-23_ was from Phoenix Pharmaceuticals (#005-26). 3-Isobutyl-1-methylxanthine (IBMX) was from Acros Organics (#228420050). PF-04447943 was purchased from MedChem Express (#HY-15441/CS-0942). Polyethylenimine (PEI) was from Polysciences, Inc. (#23966). Sodium fluoride was from EMD Chemicals (#SX0550-1). Sodium Vanadate was from Fisher Scientific (#S454). From Roche, PhosSTOP™ (#04906837001) and cOmplete™, Mini Protease Inhibitor Cocktail (#04693124001) were purchased. Protein concentration was measured using Micro BCA™ Protein Assay Kit from Thermo Scientific (#23235). cGMP was measured using Cyclic GMP XP® Assay Kit from Cell Signaling (#4360).

For Western Blotting the following primary antibodies were used: Flag (Sigma #F1804), NPRA (Novus Biologicals #NBP1-31333), NPRB (Proteintech #55113-1-AP), NPRC (Novus Biologicals #NBP1-31365), NanoLuc® Luciferase (R&D Systems #MAB10026), β-actin (Cell Signaling #4967). Secondary antibodies were goat anti-Mouse IgG−Alkaline Phosphatase (Sigma #A3562) and goat anti-Rabbit IgG–Alkaline Phosphatase (Sigma #3687). For immunoprecipitation, Anti-FLAG® M2 Affinity gel (Sigma #A2220) was used.

### Cell Culture and Transfection

HEK-293T, HEK-293FT, HeLa and A-10 cells were obtained from ATCC. HIB-1B cells (63) were a generous gift from Dr. Bruce Spiegelman. All cells were cultured with DMEM high glucose containing 10% FBS, 50 units/ml penicillin, and 50 µg/ml streptomycin. HAP-1 cells were maintained in Iscove’s modified Dulbecco’s medium with 10% FBS and 50 units/ml penicillin and 50 µg/ml streptomycin. HIB-1B cells were differentiated by 48-hour treatment with 1 µm rosiglitazone. For plasma membranes, 8 µg total plasmid was transfected using 21 µg of PEI per 10 cm plate. For co-immunoprecipitation experiments, 2 µg total plasmid was transfected using 6 µg of PEI per 3.5 cm diameter well. For luciferase assay, 0.4 µg total plasmid was transfected using 1.2 µg of PEI per well of a 96-well plate. Cells were collected twenty-four hours post transfection.

### SDS-PAGE and Western Blotting

For all Western Blots, protein was resolved in 10% Tris-glycine gels, transferred to nitrocellulose membranes, which were blocked in 5% milk for 1-hour at room temperature, then incubated overnight at 4 °C with specific primary antibodies, followed by secondary antibody incubations for 1-hour at room temperature. Western Blot image acquisition was performed on The Digital ChemiDoc MP (Bio-Rad) at the VUMC Molecular Cell Biology Resource Core, except Figure 1D was imaged on a Typhoon FLA9000 variable mode imager.

### Animals and Tissue Membrane Enriched Fractions

All animal studies were conducted at Vanderbilt University Medical Center (VUMC) and were approved by the Institutional Animal Care and Use Committee of Vanderbilt University Medical Center and in accordance with the NIH Guide for the Care and Use of Laboratory Animals. All mice used in these studies are on a C57BL/6J background. NPRC^-/-^ adipose tissue was from mice with a floxed *Npr3* allele obtained from the Knockout Mouse Project at the University of California, Davis that had been crossed to *Adiponectin-Cre* mice (JAX-010803), which we have described previously (43). At eight weeks of age mice were placed on a high-fat diet (HFD; 60% of calories from fat, Research Diet, D12492) for 12 weeks. After the 12-week feeding period, mice were euthanized, and tissues were carefully dissected, and immediately frozen in liquid nitrogen.

Membrane enriched fraction preparation was based on a previously published protocol (64). Briefly, 50 mg of tissue was homogenized in 500 µl Sucrose Buffer (250 mM sucrose, 20 mM imidazole, pH 7.0) using a TissueLyser II (Qiagen). Tissues from 3-4 mice were pooled together, triturated through a 25-guage needle, and centrifuged at 10,000×g to obtain a pellet containing nuclei, mitochondria, and large cell fragments. The supernatant is the cytosolic fraction. The pellet was then washed in 500 µl Sucrose Buffer. The pellet was resuspended in 35 µl Solubilization Buffer (50 mM NaCl, 50 mM imidazole, 2 mM 6-aminohexanoic acid, 1 mM EDTA, pH 7.0) by triturating. Twenty µl of 20% digitonin was added and the mixture was incubated at room temperature 5-10 minutes. The mixture was centrifuged at 20,000×g for 15 minutes and 40 µg protein from the supernatant was used for Western Blotting.

### Cell Lysate and Immunoprecipitation

Cells were lysed and sonicated in a buffer containing 25 mM HEPES (pH 7.4), 150 mM NaCl, 5 mM EDTA, 5 mM EGTA, 5 mM glycerophosphate, 0.9% Triton X-100, 0.1% IGEPAL, 5 mM sodium pyrophosphate, 10% glycerol, plus 1 tablet each of PhosSTOP™ and cOmplete™, Mini Protease Inhibitor Cocktail per 10 ml of lysis buffer. For immunoprecipitations, 1 mg total protein from the cell lysate was incubated with Anti-FLAG® M2 Affinity gel at 4 °C for 2 hours, followed by three 5 min washes with the lysis buffer. Immunoprecipitate or 50 µg total protein from lysate were used for Western Blotting.

### NanoLuc Luminescence Assay

NanoLuc® Binary Technology (NanoBiT) is a two-subunit luciferase system used for detection of protein:protein interactions. Following the NanoBiT® PPI MCS Starter System protocols, plasmids containing Large BiT (LgBiT; 17.6kDa, 156 amino acids) and Small BiT (SmBiT; 11 amino acids) subunits were fused to the C-terminus of full length and truncated NPRA, NPRB, NPRC (**Supplemental Figure 1**, **Supplemental Table 1**). In addition, positive control vectors containing the PKA regulatory (LgBiT-PRKAR2A) and catalytic (SmBiT-PRKACA) subunits were utilized for protocol validation (data not shown). Equimolar ratios of LgBiT and SmBiT plasmids were transiently co-transfected into HAP-1 cells growing in flat, clear-bottom, black-wall 96-well plates (Greiner Bio-One #655090). Thirty-six hours post transfection, protein:protein interactions were detected in real-time using live cells with the Nano-Glo® Live Cell Assay System, a nonlytic detection reagent that contains the cell-permeable substrate furimazine. Experiments were performed on a Synergy™ NEO HTS Multi-Mode Microplate Reader (Biotek Instruments) in the Vanderbilt High-Throughput Screening (HTS) Core Facility. For each experiment, conditions were run in triplicate. Certain replicates were excluded if contaminated by light from neighboring wells. Data are from four independent experiments. Data were log transformed and analyzed by paired t-test comparing each condition to the untransfected control using Prism 8.0.

### Duolink Proximity Ligation Assay

Duolink proximity ligation assays were performed using a Duolink™ In Situ PLA Kit Goat/Rabbit (Olink Biosciences) purchased from Sigma. Briefly, HeLa and A-10 cells were grown on 8-well glass chamber slides overnight, washed three times with cold PBS, and fixed for 10 min with 4% PFA. The chamber was removed, washed once more with cold PBS, then incubated in 70 ml 300 mM glycine PBS-T for 20 min, and one final PBS wash for 10 min. The antibodies used in the Duolink assays were rabbit Anti-NPR-A antibody from Abcam (#ab14356), at 1:200 dilution; and the goat Anti-NPR-C Antibody (N-20) from Santa Cruz Biotechnology (#sc-16871) at 1:100 dilution. Forty µl primary antibody solution was used for each well and was incubated in a humidified chamber at 4°C overnight. Secondary antibodies conjugated with the PLA probes provided by Duolink were added, and the remainder of the protocol was as described by the manufacturer. Fluorescence was detected by Nikon Eclipse-Ti TIRF microscope with Element software.

### cGMP Production Assay

HEK-293T cells were transfected with equimolar concentrations of either hNPRA-Flag and hNPRC-Flag or hNPRA-Flag and pcDNA3-Clover plasmids. At the time of harvesting, cells were rinsed with ice-cold Membrane Lysis Buffer (15 mM HEPES, 5.0 mM EDTA, 5.0 mM EGTA, 1.5 mM MgCl_2_, 2 mM PMSF, pH 7.4-7.6). Membranes made with phosphatase inhibitors had additional 10 mM NaF and 200 µM orthovanadate throughout all steps. Orthovanadate was made fresh each day by making a mixture of 50 mM vanadate and 0.16% hydrogen peroxide. Five ml ice-cold Membrane Lysis Buffer was added to each 10 cm plate, cells were scrapped off with a rubber policeman, and were triturated through a 21-gauge needle 5 times. This lysate was gently layered onto a cushion of 60% sucrose solution in Membrane Lysis Buffer in 10 cm x 1.5 cm diameter Sorvall centrifuge tubes. Lysates were centrifuged at 35,000 ×g for one hour at 4 °C in a Sorvall SS34 rotor (Du Pont). Membranes were isolated from the sucrose-buffer interface, triturated through a 25-gauge needle 5 times, aliquoted, and stored at -80 °C. Twenty-five µg of plasma membrane was used for Western Blotting. For guanylyl cyclase reaction, membranes were diluted to a concentration of 25 µg per 10 µl in Membrane Lysis Buffer. Ten µl (25 µg) membranes were added to a tube on ice containing 90 µl of Guanylyl Cyclase Reaction Buffer (50 mM Tris-HCl pH 7.6, 4 mM MgCl_2_, 1 mM GTP, 2 mM IBMX, 300 nM PF-04447943, 1 mg/ml BSA, 7.5 mM phosphocreatine, 55 units/ml creatine phosphokinase) containing 100 nM ANP. The reaction was started when tubes were placed in a 37 °C water bath. At the indicated timepoints, 20 µl was removed and placed in 180 µl Stop Reaction Buffer (50 mM Sodium acetate, pH 6.2), boiled for 3 minutes, and subsequently incubated on ice for 15 minutes. Samples were centrifuged at 12,000 ×g for 5 minutes 4 °C and supernatant was saved and stored at -80 °C. cGMP was measured in duplicate according to manufacturer’s directions using the Cyclic GMP XP® Assay Kit (Cell Signaling #4360) after the supernatant was diluted 1:10 in 1× Cell Lysis Buffer (Cell Signaling #9803). Plates were read on an Epoch Microplate Spectrophotometer with Gen5 Software (Biotek Instruments). Results were from four independent experiments using two independent preparations of membranes. The average from each sample duplicate was plotted as percent cGMP at time 0 and the linear regression function in Prism was used to calculate and determine if the slopes of cGMP concentration over time were different between membranes from NPRA+NPRC and NPRA transfected cells.

## Author contributions

Conceptualization: (DL, RPC, SC); Methodology (DL, RPC, HF); Formal analysis (RPC); Investigation (DL, RPC, WZ, HF); Resources (SC); Writing - Original Draft (DL, RPC, SC); Funding acquisition (SC).

## Classification

Biological Sciences: Physiology

## Data and materials availability

Plasmid constructs used in heterodimer studies are available upon completion of a Material Transfer Agreement. All data is available in the main text or the supplementary materials.

## Acknowledgments

We thank Dr. Todd Gulick for his insightful suggestions that contributed to this study and Dr. Thomas Maack for valuable advice and discussions. Dr. Michaela Kuhn kindly provided the NPRC-YFP plasmid; Dr. Michael Chinkers kindly provided Flag-rNPRA plasmid; Dr. Timo Müller kindly provided Flag-p62 plasmid. We thank Drs. Joshua Bauer and Dehui Mi of the Vanderbilt Institute of Chemical Biology for assistance in developing the NanoBIT assay for high-throughput screening.

