## Supplemental for "Discovery of another mechanism for the inhibition of particulate guanylyl cyclases by the natriuretic peptide clearance receptor"

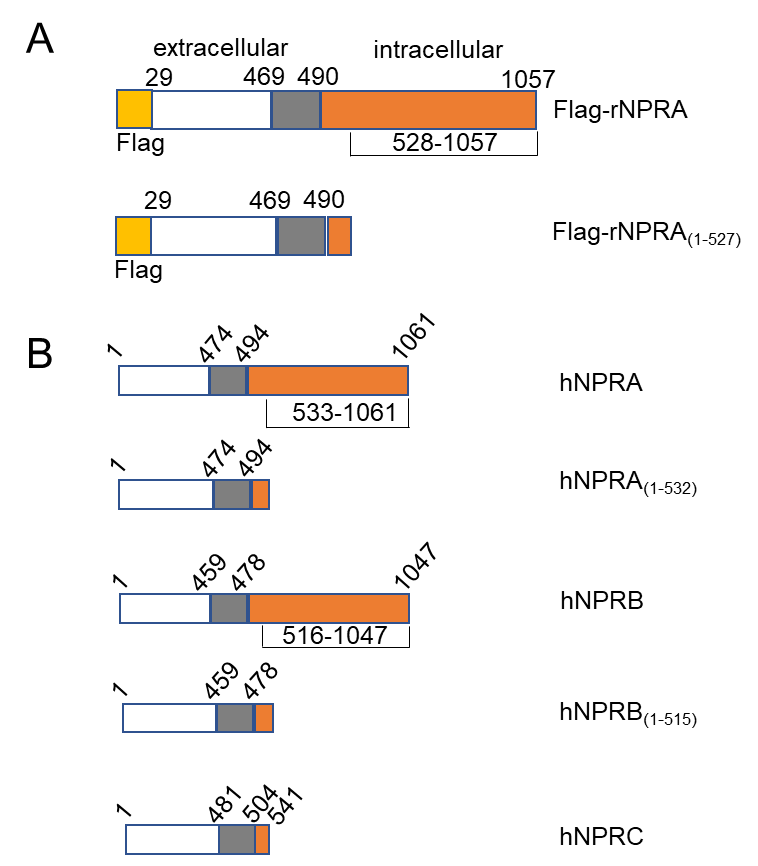

Supplemental Figure S1. Diagram of synthetic natriuretic peptide receptor constructs

The synthetic natriuretic peptide receptor constructs used in this study are indicated with the extracellular domain in white, the transmembrane region in grey, the intracellular region in orange, and any epitope tag in yellow. (**A**) Diagram of full length and truncated rat NPRA with an N-terminal Flag epitope. (**B**) Diagram of full length and truncated versions of human NPRA, NPRB, and NPRC used in these studies. Epitopes for the complementation reporter assay (LgBiT and SmBiT) were on the C-terminus.

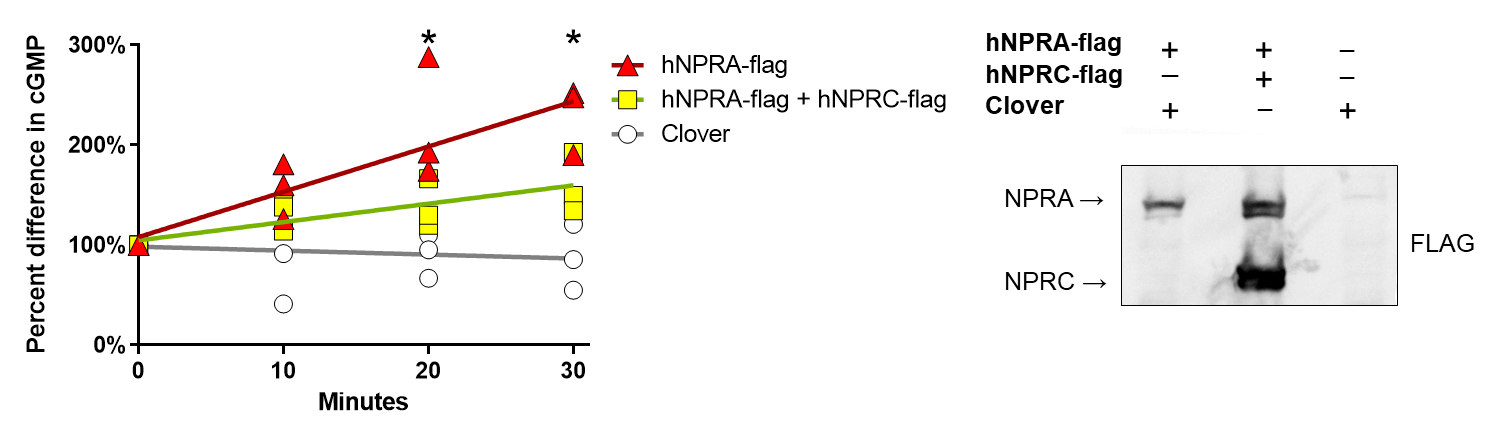
Supplemental Figure S2. Co-expression of NPRC reduces ANP-evoked cGMP production in plasma membranes

Plasma membranes were made with phosphatase inhibitors from HEK-293T cells transfected with either NPRA-flag plus pcDNA3-Clover control or NPRA-flag plus NPRC-flag. Co-expression of hNPRC along with hNPRA reduced the cGMP generated in the presence of 100 nM ANP (P=0.0203, test for difference in slopes between NPRA and NPRA+NPRC; Šídák's multiple comparisons test comparing NPRA vs. NPRA+NPRC are indicated on the graphs). Cells transfected with pcDNA3-Clover did not appear to generate cGMP. Western Blot of membranes used in the ANP-stimulated cGMP assay.

| Expressed Protein | Species | Gene | Accession No. | Vector | From | Catalog # | Reference |
| --- | --- | --- | --- | --- | --- | --- | --- |
|  |  |  |  | pcDNA3.1 | Invitrogen |  |  |
| Flag-p62 | human | SQSTM1 |  | pcDNA3 | Timo Müller |  | (65) |
| HA-VEGFR1-Myc-Flag | human | VEGFR1 |  | pCMV6 Entry | Christie Thomas | Addgene #83435 | (66) |
| GC-C-Flag | human | GUCY2C | NM_004963 | pcDNA3.1^+^/C-(K)-DYK | GenScript | OHu14859D |  |
| Flag-NPRA | rat | Npr1 |  | pCMV6 | Michael Chinkers |  | (57) |
| Flag-NPRA(1-527) | rat | Npr1 |  | pCMV6 | Dianxin Liu |  |  |
| NPRA-Flag | human | NPR1 | NM_000906 | pcDNA3.1^+^/C-(K)-DYK | GenScript | OHu09254D |  |
| NPRA-LgBiT | human | NPR1 | NM_000906 | pBiT1.1-C[TK/LgBiT] | Dianxin Liu |  |  |
| NPRA-SmBiT | human | NPR1 | NM_000906 | pBiT2.1-C[TK/SmBiT] | Dianxin Liu |  |  |
| NPRA(1-532)-LgBiT | human | NPR1 | NM_000906 | pBiT1.1-C[TK/LgBiT] | Dianxin Liu |  |  |
| NPRA(1-532)-SmBiT | human | NPR1 | NM_000906 | pBiT2.1-C[TK/SmBiT] | Dianxin Liu |  |  |
| NPRB-Flag | human | NPR2 | NM_003995 | pcDNA3.1^+^/C-(K)-DYK | GenScript | OHu18377D |  |
| NPRB(1-515)-LgBiT | human | NPR2 | NM_003995 | pBiT1.1-C[TK/LgBiT] | Dianxin Liu |  |  |
| NPRB(1-515)-SmBiT | human | NPR2 | NM_003995 | pBiT2.1-C[TK/SmBiT] | Dianxin Liu |  |  |
| NPRC-YFP |  |  |  |  | Michaela Kuhn |  |  |
| NPRC-Flag | human | NPR3 | NM_001204375 | pcDNA3.1^+^/C-(K)-DYK | GenScript | OHu09262D |  |
| NPRC-LgBiT | human | NPR3 | NM_001204375 | pBiT1.1-C[TK/LgBiT] | Dianxin Liu |  |  |
| NPRC-SmBiT | human | NPR3 | NM_001204375 | pBiT2.1-C[TK/SmBiT] | Dianxin Liu |  |  |
| LgBiT-PRKAR2A |  | PRKAR2A |  | LgBiT-PRKAR2A Control Vector | Promega | N203A |  |
| SmBiT-PRKACA |  | PRKACA |  | SmBiT-PRKACA Control Vector | Promega | N204A |  |
| Large BiT |  |  |  | pBiT1.1-C[TK/LgBiT] | Promega | N196A |  |
| Small BiT |  |  |  | pBiT2.1-C[TK/SmBiT] | Promega | N197A |  |
| Clover |  | Clover |  | pcDNA3 | Michael Lin | Addgene #40259 | (67) |

Table S1. Plasmid Table

| Primers for cloning into pBiT1.1-C[TK/LgBiT] and pBiT2.1-C[TK/SmBiT] | |
| --- | --- |
| NPR1-F | 5’-TCTGCTAGCATGCCGGGGCCCCGGCGCCCCGCTGGCTCCCGCCTGCGCCTGCTC-3’ |
| NPR1-R | 5’-CCGCTCGAGCCTCGGGTGCTACTCCCCCTCTCCCCAAG-3’ |
| NPR1-532-R | 5’-CCGCTCGAGCCCAGGGTCAGCCGGCTGCCTGCACTCCG-3’ |
| NPR2-F | 5’-CCCAGATCTATGGCGCTGCCATCACTTCTGCTGTTGGTG-3’ |
| NPR2-R | 5’-CCACTCGAGCCCAGGAGTCCAGGAGGTCCTTTCCGCT-3’ |
| NPR2-515-R | 5’-AAACTCGAGCCGAGGCGACTGCCTGCACCTTTGTGATAAC-3’ |
| NPR3-F | 5’-TCTGCTAGCATGCCGTCTCTGCTGGTGCTCACT-3’ |
| NPR3-R | 5’-CCGCTCGAGCCAGCTACTGAAAAATGGGATCTGATGG-3’ |
| Primers for site-directed mutagenesis | |
|  | 5’-GCTGGCAGCCTTCTGTAGACCCTGAGTGGGCGA-3’ |
|  | 5’-TCGCCCACTCAGGGTCTACAGCCGGCTGCCAGC-3’ |
| Primers for sequencing | |
| BiT Forward | 5’-AAGCTTGGCAATCCGGTACT-3’ |
| BiT lg Reverse | 5’-GTTCCCAGTCCCCAACGAAATC-3’ |
| Sm BiT Reverse | 5’-CAGAATCTCCTCGAACAGCCGGTAGCCGGTCAC-3 |

Table S2. Primers for cloning and sequencing confirmation
